# What’s in your next-generation sequence data? An exploration of unmapped DNA and RNA sequence reads from the bovine reference individual

**DOI:** 10.1101/022731

**Authors:** Lynsey K. Whitacre, Polyana C. Tizioto, JaeWoo Kim, Tad S. Sonstegard, Steven G. Schroeder, Leeson J. Alexander, Juan F. Medrano, Robert D. Schnabel, Jeremy F. Taylor, Jared E. Decker

## Abstract

Next-generation sequencing projects commonly commence by aligning reads to a reference genome assembly. While improvements in alignment algorithms and computational hardware have greatly enhanced the efficiency and accuracy of alignments, a significant percentage of reads often remain unmapped. We generated *de novo* assemblies of unmapped reads from the DNA and RNA sequencing of the *Bos taurus* reference individual and identified the closest matching sequence to each contig by alignment to the NCBI non-redundant nucleotide database using BLAST. As expected, many of these contigs represent vertebrate sequence that is absent, incomplete, or misassembled in the UMD3.1 reference assembly. However, numerous additional contigs represent invertebrate species. Most prominent were several species of Spirurid nematodes and a blood-borne parasite, *Babesia bigemina*. These species are not known to infect taurine cattle and the reference animal appears to have been host to unsequenced sister species. We demonstrate the importance of exploring unmapped reads to ascertain sequences that are either absent or misassembled in the reference assembly and for detecting sequences indicative of infectious or symbiotic organisms.

## Introduction

Next-generation sequencing has vastly increased the dimensionality of sequencing projects and routinely allows the generation of hundreds of millions or even billions of short reads. Analysis of these data requires that the short reads be assembled into contiguous sequences either using *de novo* or reference-guided assembly. For organisms with a reference genome, reads generated in the sequencing process are usually matched to the reference sequence with a variety of alignment algorithms. This is currently the most efficient way of transforming the raw sequence reads into a consensus sequence. However, there are several limitations inherent to the alignment process, including alignment to repetitive regions, absent or misassembled sequence in the reference genome, and individual genetic divergence between the subject organism’s genome and the reference genome^1^. Despite these challenges, the majority of reads produced from a sequencing experiment will adequately align to a reference assembly. Nevertheless, a small but significant fraction of reads frequently remain unmapped.

Unmapped reads have generally been disregarded and these data are often discarded. However, recent work has begun to focus on the development of bioinformatic tools for detecting pathogens in human sequence data by the computational subtraction of known human sequences^2–4^. Application of these pipelines in other recent studies has suggested that potentially biologically relevant information can be extracted from the unmapped reads^5,6^. Using an original alignment, assembly, and identification pipeline that can be applied to data from any species, we took advantage of a unique opportunity to explore the unmapped reads from the DNA and RNA sequencing of L1 Dominette 01449, the *Bos taurus* reference individual^7^. These data had not previously been used in the creation or annotation of the reference assembly.

By using sequence data produced from the reference individual, we minimize alignment challenges that are due to genetic variation among individuals. We identified DNA and RNA contigs that were assembled *de novo* from unmapped reads that could generally be classified into one of three categories: 1) sequence from bovine; 2) sequence from other vertebrate species that was orthologous to bovine; and 3) sequence from non-vertebrate species. Our analysis unequivocally demonstrates that the unmapped reads contain important data pertaining to sequences from the organism that are missing from the reference assembly, represented by categories 1 and 2, and sequences that can be used to identify microbiota members, putatively represented by category 3.

## Results

### De novo assembly of unmapped reads

Approximately 111.7 million DNA sequence reads, 7.2% of the total, remained unmapped after alignment to the reference genome. A fraction of those reads could be used for assembly, due to a large number of sequences with low quality (Table S1). However, approximately 1.4 million reads were incorporated into 69,230 contigs with an N50 of 737 bp. Overall, the contigs comprised approximately 46.6 Mb. Additional assembly statistics are provided in Table S1.

A median of approximately 6.7% of RNA-seq reads remained unmapped for each of the 17 tissue samples. *De novo* assembly of these reads yielded a total of 43,961 contigs, with a median of 1,792 contigs per tissue and an N50 of 324.5 bp. Overall, the contigs spanned 14.8 Mb with a median of 603 Kb per tissue. Assembly statistics for each tissue are in Table S2.

### Pairwise alignment of contigs assembled from unmapped DNA reads to the non-redundant nucleotide database

Approximately 51% of the contigs generated from the unmapped DNA reads produced a significant alignment when queried against the non-redundant nucleotide database (*nt*) database. The most common alignment was to other *Bos taurus* sequences (Fig. 1). This result was expected given the draft quality of the bovine reference assembly and considering that we assembled paired reads if either one or both of the reads were unmapped to the reference assembly. However, the second most common alignment for these DNA contigs was to *Onchocerca ochengi*, a nematode known to infect indicine cattle that has been heavily researched due to its similarity to the parasite that causes African River Blindness in humans. We simulated paired-end sequence read data from the *O. ochengi* genome assembly by randomly shearing the genome and then aligned the produced paired-end reads to the bovine reference assembly and concluded that the *O. ochengi* assembly, based on samples obtained from cattle skin^8^, is contaminated with cow sequences.

**Figure 1.**
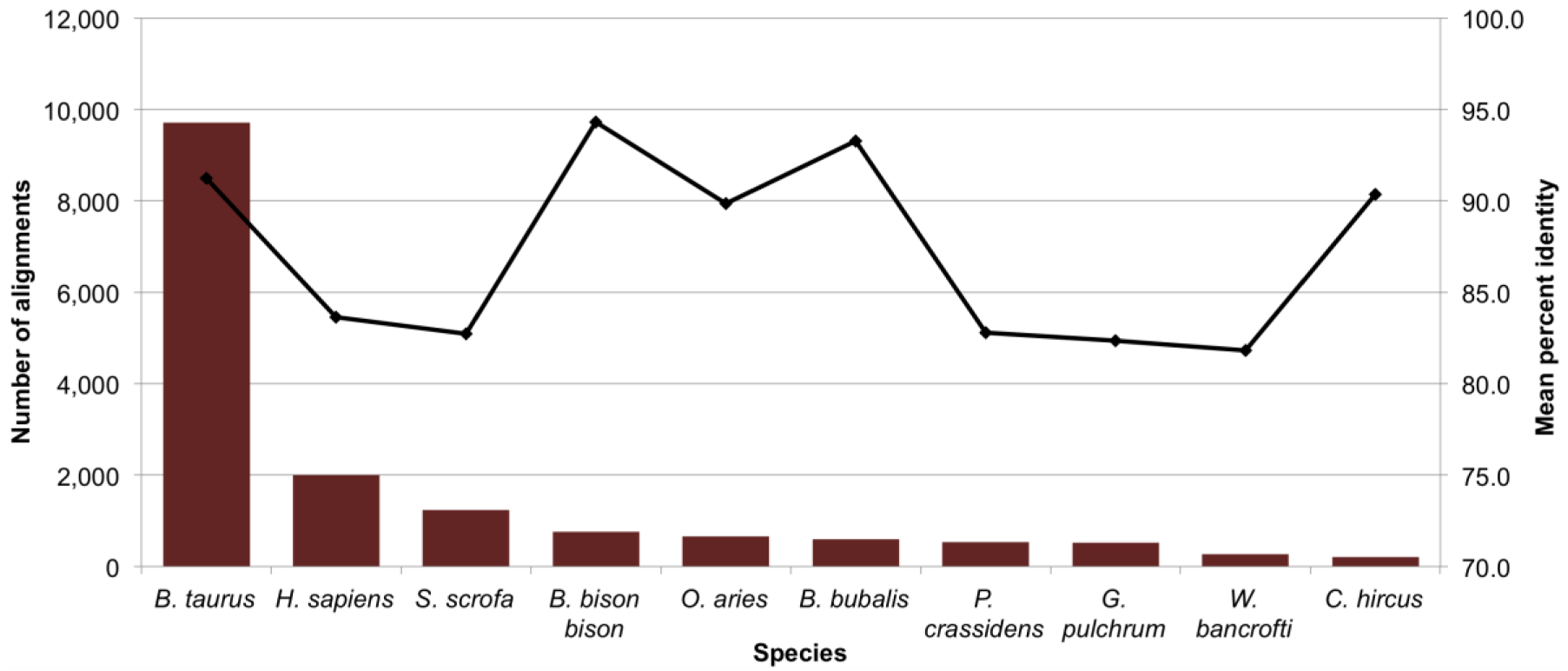
Most common alignments from DNA. Ten most common species with significant alignments from the pairwise alignment of *de novo* assembled contigs from unmapped DNA reads and their mean percent identity.

Consequently, we excluded the *O. ochengi* assembly from any further analyses (Supplementary Note 1).

On subsequent analyses preventing alignment to *O. ochengi*, a fraction of the contigs originally identified as *O. ochengi* were unambiguously matched to bovine sequences. However, the number of alignments to other filarial nematode sequences also increased drastically. These included hundreds of contigs aligned to *Gonglyonema pulchrum* and *Wuchereria bancrofti,* and a few to *Parascaris equorum. G. pulchrum* and *W. bancrofti* belong to the order Spirurida, as does *O. ochengi*, but are known to only infect humans. The alignments to each of these species had a percent identity of approximately 82% (Table 1), which is consistent with cattle not being a host of these nematodes and indicates that these represent an alignment to a previously unsequenced sister species of *G. pulchrum* and *W. bancrofti*.

**Table 1.** Top four non-vertebrate alignments to *de novo* assembled contigs from unmapped DNA sequence reads.

| Species | Number of Alignments | Mean Identity (%) | Maximum Identity (%) | Mean Length (bp) | Maximum Length (bp) | Mean E-Value |
| --- | --- | --- | --- | --- | --- | --- |
| *Gongylonema pulchrum* | 516 | 82.34 | 100 | 603.97 | 1,008 | 1.78E-05 |
| *Wuchereria bancrofti* | 273 | 81.81 | 98.82 | 742.81 | 1,607 | 3.65E-28 |
| *Babesia bigemina* | 190 | 91.23 | 100.00 | 502.47 | 2,078 | 1.35E-05 |
| *Parascaris equorum* | 68 | 88.49 | 100 | 566.53 | 1,008 | 1.31E-04 |

**Table 2.** Top four non-vertebrate alignments to *de novo* assembled contigs from unmapped RNA-seq reads.

| Species | Number of Alignments | Mean Identity (%) | Maximum Identity (%) | Mean Length (bp) | Maximum Length (bp) | Mean E-Value |
| --- | --- | --- | --- | --- | --- | --- |
| *Uncultured bacterium* | 34 | 96.70 | 100.00 | 355.44 | 1,381 | 2.35E-83 |
| *Bovine herpesvirus 6* | 22 | 99.09 | 100.00 | 462.68 | 933 | 2.73E-100 |
| *Onchocerca flexuosa* | 13 | 87.58 | 89.12 | 253.54 | 510 | 8.46E-42 |
| *Babesia bigemina* | 12 | 93.46 | 99.51 | 390.50 | 926 | 5.00E-69 |

Also detected was *Babesia bigemina,* a blood-borne parasite known to cause bovine babesiosis, or Texas fever in cattle. While only 190 contigs aligned to *B. bigemina*, significantly less than the combined number of alignments to nematode species, ten were larger than 1,000 bp and the mean identity was 91.13% (Table 1). Consequently, we hypothesize that an unsequenced sister specie of *B. bigemina* had infected Dominette and was detected via sequence. A complete summary of significant alignments, both vertebrate and non-vertebrate, is presented in Table S3.

### Pairwise alignment of contigs assembled from unmapped RNA-seq reads to the non-redundant nucleotide database

The pairwise alignment of the *de novo* assembled contigs generated from the unmapped RNA-seq reads to the *nt* database produced similar results to the alignment of the DNA contigs. Overall, 83% of the RNA-seq contigs had significant alignments to sequences in the *nt* database. Across all tissues, *Bos taurus* was the most common alignment. Also prominent are alignments to *Bison bison bison, Bubalus bubalis,* and *Bos mutus*, all closely related to cow (Fig. 2). Significant BLAST alignments of the unmapped RNA-seq unmapped read contigs to cow or these other closely related species indicates the existence of coding regions that are missing or misassembled in the reference assembly. By mapping the GI number of the most significant BLAST alignment to a gene symbol, we detected alignments to 4,412 *B. taurus* and 4,029 *B. bison bison, B. bubalis*, or *B. mutus* genes, suggesting that as much as 42% of the bovine protein coding genome is misassembled. Further results and discussion of these analyses are included in Supplementary Note 2.

**Figure 2.**
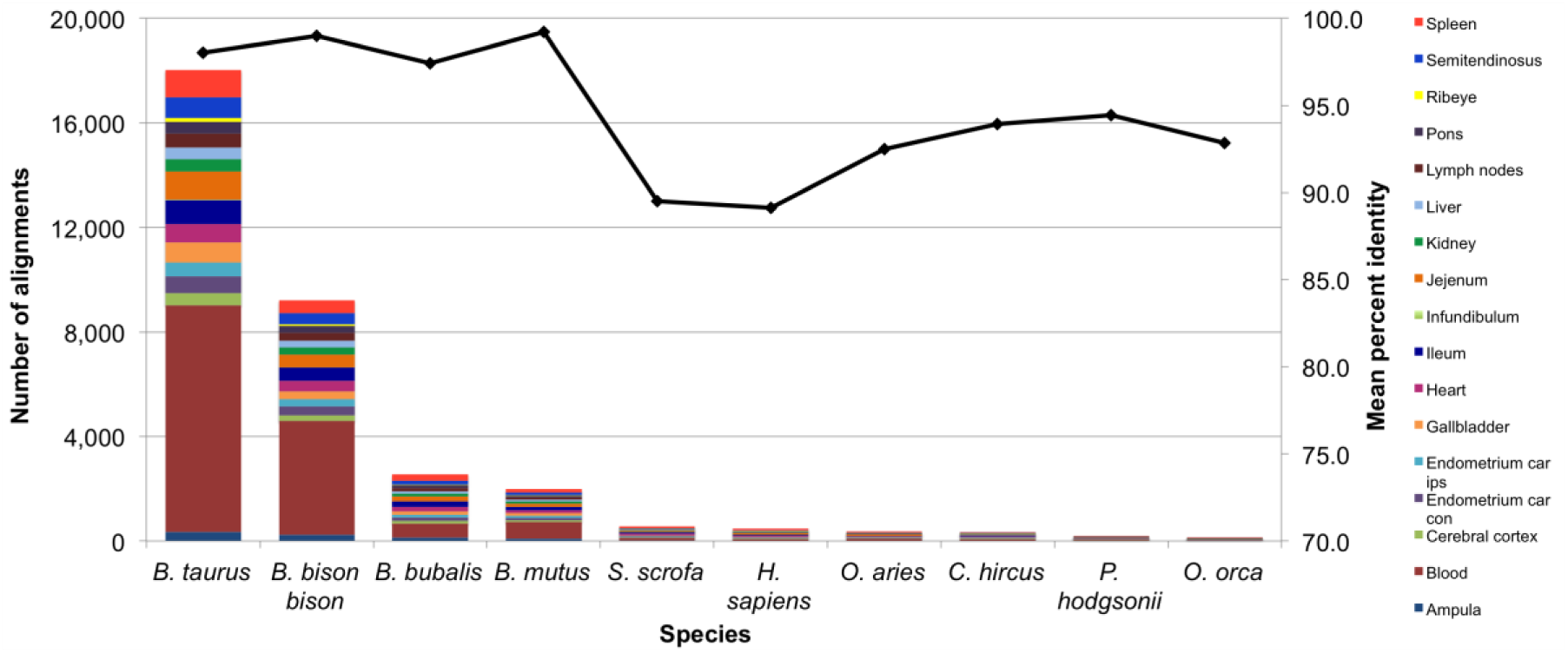
Most common alignments from RNA. Ten most common species with significant alignments from the pairwise alignment of *de novo* assembled contigs from unmapped RNA-seq reads by tissue and their overall mean percent identity.

As was the case in the pairwise alignment of DNA unmapped contigs, there were also numerous alignments to other vertebrate and non-vertebrate species. The most common alignments to non-vertebrate species included uncultured bacterium, bovine herpesvirus 6, *Onchocerca flexuosa* and *B. bigemina*. Bovine herpesvirus 6 was previously discovered as a contaminant in the UMD3.1 build by Merchant *et al*.^9^, who concluded that Dominette must have been host to the virus. Alignments to *O. flexuosa* and *B. bigemina* support the hypothesis generated from the analysis of the unmapped DNA read contigs that Dominette was also host to a nematode of the Spirurida order and an unsequenced relative of *B. bigemina*. Several additional fungal and bacterial species were also detected in the unmapped read RNA-seq contigs at low levels. A complete summary of alignments from all 17 tissues is presented in Tables S4 and S5.

## Discussion

To our knowledge, this is the first formal investigation into the nature and identity of unmapped reads from the resequencing of an individual used for the generation of a reference genome assembly. These data afforded us several luxuries such as the direct comparison of reads to the reference assembly without alignment challenges due to genetic variation between the reference and the resequenced genomes. Second, the opportunity to compare independently generated datasets from the same individual provided unequivocal support for our discovery of concordant non-vertebrate sequences within the whole genome and transcriptome sequences of the bovine host. In addition to our sequencing of cDNA generated from RNA isolated from 17 tissues, we also sequenced genomic DNA that had been isolated from both liver and white blood cells at three separate facilities. Endogenous contaminants were detected in the reads that were generated from all three sequencing runs. Nearly all of the contigs assembled *de novo* from unmapped reads that were identified as representing pathogens or symbionts comprised reads originating from sequencing libraries generated at all three facilities. These attributes facilitated both the discovery and validation of the pathogen sequences found in this study.

Despite the continuing exponential increases in sequences submitted to NCBI’s databases, the number of represented species still comprises only a small proportion of existing species. While we detected several sequence alignments to spirurid nematodes in both the DNA and RNA sequence data, none of these species are known to infect taurine cattle. Therefore, we postulate that the actually species present within the tissues of Dominette is either an undiscovered species or a previously recognized, but unsequenced, organism such as *Onchocerca gutturosa* or *Onchocerca lienalis* from the Spirurida order. Both *O. gutturosa* and *O. lienalis* are known to infect taurine cattle in various parts of the United States^10^. However, these species have not been sequenced other than for a few select genes to generate data for phylogenetic analyses^11–19^. In this study, we assembled nearly 1,000 contigs that we believe represent novel sequence from a Spirurid nematode that infects taurine cattle in North America.

The precise identity of the sequence matching *B. bigemina* in both the RNA-seq and genomic DNA data is also ambiguous. As no fever like symptoms were reported in this cow who spent her life at a USDA research facility near Miles City, Montana and babesiosis has been reported to have been eradicated in the United States with vaccination no longer being required^20^, we suspect that Dominette was asymptomatically infected with a non-pathogenic strain of *Babesia* spp., as has been previously reported in Turkey^21^, Syria^22^, and Thailand^23^. Although it is currently not possible to determine the exact species of parasite, we can assess the animal’s parasite burden via deep sequence data by evaluating the number of species to which the contigs of unmapped reads align and the number of contigs that align to each species. Parasite burden negatively impacts animal health and profitability^24,25^ and can serve as a reservoir for later infections^22^. Although symptoms were not visible and the animal appeared healthy, the detection of subclinical parasite burden is important because a physiological response to the infection must still occur. This response reduces fitness, causes a decrease in production traits such as feed intake and feed efficiency^25,26^ and can also influence the interpretation of RNA-seq experiments.

In conclusion, we alert researchers that many sequences of interest may be found in the reads that fail to align to a reference assembly. We demonstrate that the unmapped reads contain biologically significant information relative to genes that are either partially or completely missing from the reference assembly, as well as information regarding the identity and magnitude of endosymbionts or pathogens infecting the individual. The large number of missing or misassembled bovine protein coding genes must significantly impact the interpretation of RNA-seq studies and warrants further research. Continuation of unmapped read mining will also expand our knowledge of the extent of internal parasitic infections and may lead to the discovery of previously unknown symbiotic relationships. These metagenomic inferences are an additional source of information from whole-genome sequencing data that can be used as phenotypes or covariates in downstream analyses. As the quality of reference assemblies improves and the scope of sequenced microorganisms broadens, the detection of parasitic infections and symbiotic relationships will become more explicit.

## Methods

### DNA and RNA sequencing

DNA was extracted from liver and whole blood samples from L1 Dominette 01449 (referred to here as “Dominette”), a Hereford cow used to generate the *Bos taurus* Sanger reference assembly^7^, and was sent to three separate facilities for sequencing. Twenty paired-end and 12 mate-pair libraries were constructed and DNA was 2×100 bp sequenced using an Illumina HiSeq2000 to an average coverage of approximately 55X.

RNA was extracted using Trizol Reagent (Invitrogen, Carlsbad, CA) as described elsewhere^27^ from 17 tissue samples including ampula, blood, cerebral cortex, endometrium sampled from caruncular regions contralateral (car con) and ipsilateral (car ips) to the corpeus luteum, gallbladder, heart, ileum, infundibulum, jejunum, kidney, liver, mesenteric lymph nodes, pons, ribeye muscle, semitendinosus muscle, and spleen. Preparation of the mRNA samples for sequencing was performed by Global Biologics (Columbia, MO) using the TruSeq Stranded mRNA Library Prep Kit (Illumina®, San Diego, CA) and sequenced 2×100 bp, with the exception of blood which used the TruSeq RNA Sample Preparation Kit and was sequenced 1×100 bp.

### Pre-processing and alignment of reads

Error correction was performed on DNA sequence reads using the QuorUM error correction algorithm^28^. After filtering duplicate and low quality reads, 1,622,097,087 unique reads remained. Paired reads were aligned to the UMD3.1 cow reference assembly using NextGENe 2.4.1 (SoftGenetics, LLC, State College, PA) requiring at least 35 contiguous bases with ≥95.0% overall match, up to 2 allowable mismatched bases, and up to 100 allowable alignments of equal probability genome-wide.

RNA sequence reads were filtered for quality and adapter sequences were trimmed using a custom Perl script already described^27^. Computations were performed on the HPC resources at the University of Missouri Bioinformatics Consortium (UMBC). TopHat v2.0.6^29^ was used to map the reads to the *Bos taurus* UMD3.1 reference genome. A total of 2 mismatches and up to 3 bp indels were allowed in alignment.

### De novo assembly of unmapped reads

Reads from DNA sequencing that remained unmapped following alignment to the reference genome were assembled using MaSuRCA 2.3.2^30^. Reads from RNA sequencing that remained unmapped following alignment were assembled using Trinity version r20140717^31^. To maintain a paired read file structure, reads where both the forward and reverse read were unmapped or where one of the reads was unmapped but the other was mapped were collectively used for assembly.

### Pairwise alignment of unmapped contigs to the nt database

Prior to pairwise alignment, contigs assembled from the unmapped DNA reads were sorted by size and only contigs greater than 500 bases were aligned (n = 42,086). Due to the smaller size of the RNA contigs, they were not filtered by size prior to pairwise alignment. Using the blastn algorithm of BLAST+ 2.2.30^32,33^, each DNA and RNA contig was aligned to the NCBI non-redundant nucleotide database and the most significant alignment was returned. The BLAST output was then parsed to determine the subject species, percent identity, length of match, number of mismatches, number of gaps, E-value, and overall score. Significant alignments were declared only if the length of the alignment was ≥200 bp for DNA or ≥50 bp for RNA. Only the best match for each aligned contig was reported. This output was summarized according to the total number of alignments per species, average and maximum percent identity, average and maximum length of match, and average e-value.

### Quantification and identification of coding regions within unmapped reads

Contigs from unmapped RNA-seq reads were aligned to contigs from unmapped DNA reads using NextGENe 2.4.1 requiring ≥98% overall match to declare a match. Additionally, for the significant RNA alignments, the gene symbol corresponding to the GI accession number for the alignment was captured where possible and recorded using the db2db tool in bioDBnet^34^. A unique list of gene symbols was constructed and the number of significant alignments to each gene was tallied.

### Accession Codes

Raw data have been deposited in the SRA with accession number XXXX.

## Acknowledgements

Funding for this study was provided in part from the bovine species coordinators of the USDA National Institute of Food and Agriculture supported NRSP-8 National Animal Genome Research Support Program and National Research Initiative Competitive Grants numbers 2011-68004-30214, 2011-68004-30367, 2012-67012-19743, 2013-68004-20364, MO-HAAS0027, and MO-MSAS0014 from the USDA National Institute of Food and Agriculture. The authors appreciate the contributions of the Beijing Genomics Institute in generating whole genome and tissue transcriptome sequence from the bovine reference animal.

## Contributions

J.F.T., J.E.D., R.D.S., and L.K.W. designed the experiments and interpreted the results of all analyses. L.K.W. built the analysis pipeline and analyzed the sequence data.

P.C.T. did alignments and *de novo* assemblies for RNA sequence data. L.J.A. collected tissues and extracted nucleic acids. J.W.K. extracted and quantitated RNA. T.S.S.,

S.G.S. and J.F.M. sequenced genomic DNA. J.F.T., J.E.D. and R.D.S. sequenced DNA and RNA. L.K.W. wrote the manuscript and J.F.T., J.E.D. and R.D.S. edited the manuscript. All authors read the final manuscript.

## Supplementary Information

Supplementary Note 1: The *Onchocerca ochengi* reference assembly is contaminated with bovine genomic sequence.

Supplementary Note 2: Estimation of the number of protein coding genes missing or misassembled in the UMD3.1 bovine reference assembly.

Supplementary Tables: Tables S1 through S7 describing assembly, alignment, and gene summaries.

